# Mechanism Underlying Anti-Markovnikov Addition in the Reaction of Pentalenene Synthase

**DOI:** 10.1101/2020.06.17.157867

**Authors:** Jason O. Matos, Ramasamy P. Kumar, Alison C. Ma, MacKenzie Patterson, Isaac J. Krauss, Daniel D. Oprian

## Abstract

Most terpene synthase reactions follow Markovnikov rules for formation of high energy carbenium ion intermediates. However, there are notable exceptions. For example, pentalenene synthase (PS) undergoes an initial anti-Markovnikov cyclization reaction followed by a 1,2-hydride shift to form an intermediate humulyl cation with positive charge on the secondary carbon C9 of the farnesyl diphosphate substrate. The mechanism by which these enzymes stabilize and guide regioselectivity of secondary carbocations has not heretofore been elucidated. In an effort to better understand these reactions, we grew crystals of apo-PS, soaked them with the non-reactive substrate analog 12,13-difluorofarnesyl diphosphate, and solved the x-ray structure of the resulting complex at 2.2 Å resolution. The most striking feature of the active site structure is that C9 is positioned 3.5 Å above the center of the side chain benzene ring of residue F76, perfectly poised for stabilization of the charge through a cation-π interaction. In addition, the main chain carbonyl of I177 and neighboring intramolecular C6,C7-double bond are positioned to stabilize the carbocation by interaction with the face opposite that of F76. Mutagenesis experiments also support a role for residue 76 in cation-π interactions. Most interesting is the F76W mutant which gives a mixture of products that likely result from stabilizing a positive charge on the adjacent secondary carbon C10 in addition to C9 as in the wild-type enzyme. The crystal structure of the F76W mutant clearly shows carbons C9 and C10 centered above the fused benzene and pyrrole rings of the indole side chain, respectively, such that a carbocation at either position could be stabilized in this complex, and two anti-Markovnikov products, pentalenene and humulene, are formed. Finally, we show that there is a rough correlation (although not absolute) of an aromatic side chain (F or Y) at position 76 in related terpene synthases from *Streptomyces* that catalyze similar anti-Markovnikov addition reactions.

---

Terpenes comprise the largest class of natural products with over 80,000 unique members identified to date.^*4*^ They are found largely in the plant world, but also serve important roles in animal and bacterial metabolism. They are important commercially and medicinally, and increasingly viewed as viable candidates for renewable biofuels.

The committed step in the biosynthesis of all terpenes is catalyzed by a terpene synthase (or terpene cyclase).^*5*^ Typically, these remarkable enzymes take a linear, prenyl substrate and convert it into a cyclic hydrocarbon by generating and controlling the reactivity of high energy carbenium ion intermediates in reactions that often involve complicated carbon skeleton rearrangements, including methyl and hydride migrations. Class 1 terpene synthases employ prenyl diphosphate substrates and use metal ions to trigger dissociation of the allylic diphosphate to form an initial resonance-stabilized carbenium ion intermediate.^*5*^ Cyclization follows as a result of addition reactions involving π electrons from double bonds present in the prenyl chain.^*6, 7*^ The reactions are typically terminated by deprotonation to form an olefin or cation capture by a nucleophile in the active site such as a water molecule. Downstream enzymes, mostly P450s and acyl transferases, then modify the hydrocarbon scaffold to increase functionalization of the resulting end product in the pathway.

Pentalenene synthase (PS) is a class 1 sesquiterpene synthase that catalyzes the committed step in biosynthesis of pentalenolactone antibiotics in several species of *Streptomycetes*. ^*8, 9*^ The x-ray crystal structure of the enzyme from *Streptomyces UC5319* was first solved by Christianson and coworkers in 1997.^*10*^ It is a 38 kDa monomer composed of a single domain with typical class I a-helical terpenoid cyclase fold and contains conserved Asprich (DDXXD) and NSE/DTE motifs for binding of the diphosphate moiety of the substrate along with three Mg^2+^ ions.^*5, 10*^ The Mg^2+^-dependent reaction mechanism has been studied extensively^*17–13*^ and begins with anti-Markovnikov attack of the 10,11 double bond on C1 of the farnesyl diphosphate substrate by either direct displacement of the diphosphate or through ionization and intermediary formation of a resonance-stabilized allylic carbenium ion on C1 (Fig. 1). This step in the reaction initially produces humulyl cation A, with positive charge located on the secondary carbon C10, and then humulyl cation B, following a 1,2-hydride shift to move the charge to C9. Concerted, asynchronous attack of the C2,C3 and C6,C7 alkenes leads to the protoilludyl cation intermediate,^*14–16*^ which is thought then to undergo rearrangement to the Z-secoilludyl cation, followed by a 1,2-hydride shift, transannular cyclization, and finally deprotonation to produce pentalenene (Fig. 1). This series of reaction steps produces a stereochemically dense, enantiomerically pure triquinane structure containing four chiral carbons.

**Fig. 1.**
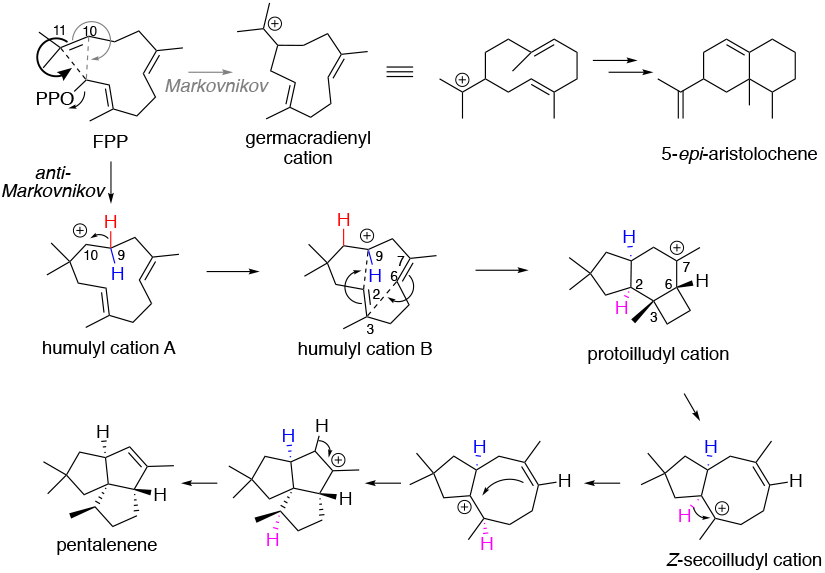
Comparison of reaction intermediates for two sesquiterpene synthases: 5-epi-aristolochene synthase and pentalenene synthase. Both reactions begin with ionization or direct displacement of the allylic diphosphate substrate (FPP) with attack of the 10,11 double bond. 5-epi-aristolochene synthase favors attack of the π electrons to form the expected Markovnikov product with positive charge on the more highly substituted carbon (C11 of the germacradienyl cation). In contrast, pentalenene synthase directs anti-Markovnikov attack such that the positive charge develops on the less substituted secondary carbon (C10) to produce first humulyl cation A, and then, following a 1,2-hydride shift, humulyl cation B with carbocation at position C9.

Our interest in this reaction is focused on how the enzyme directs anti-Markovnikov attack of C11 on C1 to produce first humulyl cation A, with the carbenium ion located on the secondary carbon C10, and then, following the 1,2-hydride shift, humulyl cation B, with the carbenium ion located on the secondary carbon at C9. How is it that the enzyme selects a pathway with the less stable secondary carbocations rather than allowing Markovnikov attack of C10 on C1 to produce the much more stable tertiary cation at C11, as for example happens in solution and with most of the other sesquiterpene synthase reactions (e.g., 5-*epi*-aristolochene synthase in Fig. 1)?^*17–20*^ How does the enzyme stabilize charge on these secondary carbon atoms? Despite the fact that pentalenene synthase was one of the first terpene cyclases to have its structure determined by x-ray crystallography,^*10*^ the enzymatic mechanism underlying regioselectivity in formation of these unstable carbenium ion intermediates on secondary carbon atoms is completely unknown.

## RESULTS

To explore the mechanism underlying the initial anti-Markovnikov cyclization in the PS reaction, we first determined a structure for the enzyme with ligand in the active site. While the structure of apo-PS was determined more than 20 years ago, a structure with ligand bound in the active site has not previously been reported. We began by growing crystals of the apo-enzyme. The crystals were then soaked with the non-reactive substrate analog 12,13-difluorofarnesyl diphosphate (DFFPP), which we prepared by enzymatic elongation of 8,9-difluorogeranyl diphosphate (DFGPP)^*21, 22*^ with isopentenyl diphosphate (IPP; Fig. 2). The two electronegative fluorine atoms in DFFPP are positioned to inhibit reactivity of the π electrons in the C10,C11-double bond and thus block the initial cyclization reaction. After 1 hr incubation, the crystals were immersed into liquid nitrogen, and the structure of the resulting complex was solved by x-ray diffraction in space group *P63* to 2.2 Å resolution.

**Figure 2.**
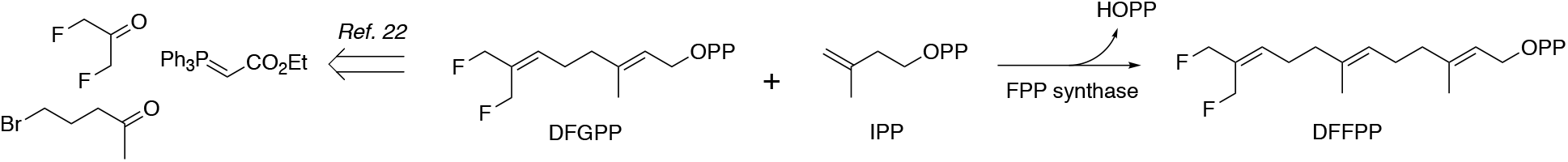
Enzymatic reaction scheme for production of 12,13-DFFPP from 8,9-DFGPP and IPP.

As has been described before for crystals of apo-PS (1PS1),^*10*^ there are two molecules in the asymmetric unit, with essentially identical secondary structure (0.3 Å RMSD). Each monomer is composed of 11 a-helices (A-K). Electron density is not visible for residues 159-165, 225-234, and 240-249 in molecule A, and these amino acids were not modelled into the structure.

As shown in Fig. 3, there is well-defined electron density in the active site of the enzyme at 1 σ contour corresponding to the inactive substrate analog DFFPP. The ligand binds only to molecule A of the asymmetric unit. As is the case with all of the structures reported here, there is an unidentified electron density in the active-site of the apo-enzyme that is displaced upon soaking with the DFFPP ligand. The diphosphate moiety of DFFPP is anchored at the conserved Asp-rich (DDXXD) and NSE/DTE motifs that are required for binding of the trinuclear metal cluster, but only two of the three Mg^2+^ ions are visible in the active site (metal ions are shown in Fig. S1). The prenyl chain extends deep into the binding pocket, makes a U-turn at the C6,C7-double bond, and then folds back on itself to position C11 4.0 Å directly across from C1 where attack of the C10,C11 π electrons on the developing allylic carbenium ion would proceed with the expected inversion of configuration at C1.^*11*^ The most striking feature of the active site structure is that C9 of the prenyl chain, the site of the reactive secondary carbocation, is positioned 3.5 Å above the center of the benzene ring of residue F76 (Fig. 4A). C10 and C8 lie clearly outside of the ring circumference. Thus, the active site architecture is organized such that C11 is located most closely to C1, and the aromatic ring of F76 is ideally positioned to stabilize a developing charge on C9, as opposed to carbons C11, C10 or C8, through a cation-π quadrupole interaction^*23–27*^ so as to support the anti-Markovnikov cyclization reaction. In addition, the developing charge on C9 is homoallylic with respect to the C6,C7-π electrons, which are also oriented plausibly for an intramolecular cation-π interaction (as described for the gas phase reaction^*14*^) with the face of C9 opposite that interacting with the F76 sidechain. Finally, the mainchain carbonyl oxygen of I177, which is exposed at the break in helix G, is oriented toward the same face of C9 as the C6,C7-double bond at a distance of 3.4 Å, well positioned to stabilize the C9 carbocation through a charge-dipole interaction (Fig. S2). The helix G break is a common feature of class 1 terpene synthases. The possible role of the helix G break in providing dipoles for stabilization of cation intermediates was first proposed by Noel and coworkers for the reaction of epi-aristolochene synthase^*28*^ and then generalized by Pandit et al.^*29*^ and Dickschat and coworkers. ^*30, 31*^

**Fig. 3.**
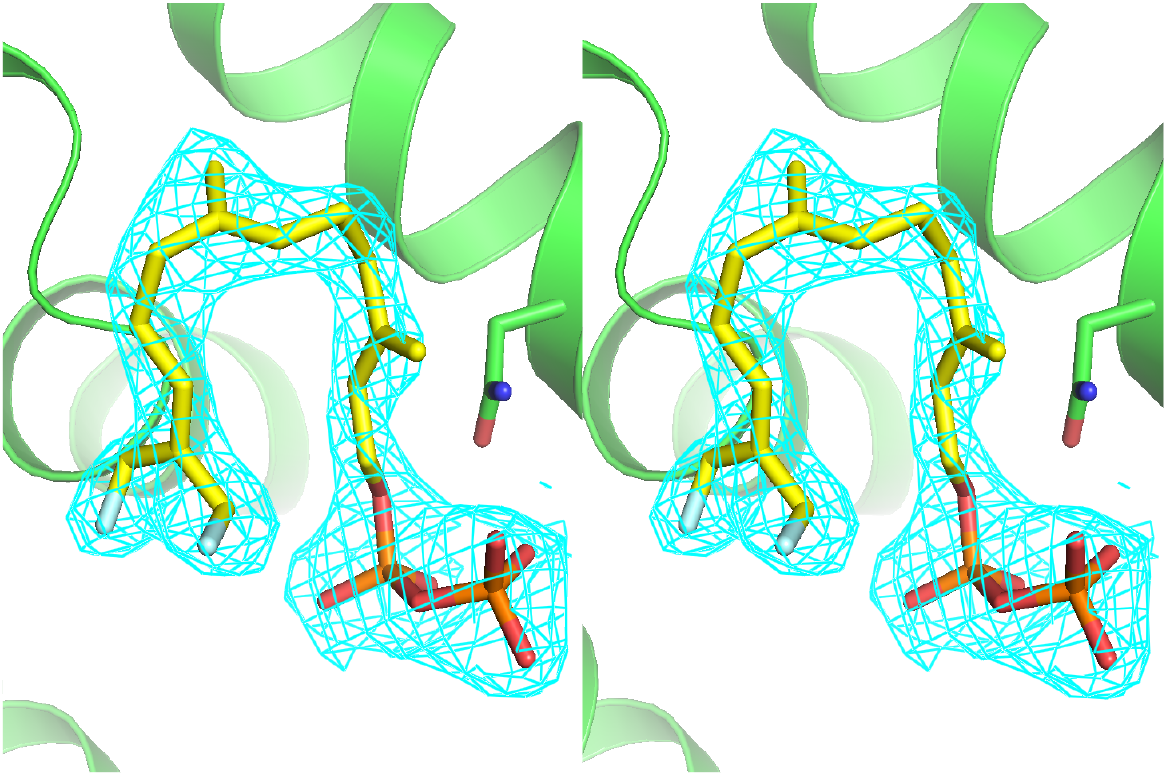
Stereo view of DFFPP ligand in the active site of PS. Fourier electron density in blue mesh is contoured at 1 σ. Protein is shown in green, diphosphate moiety of DFFPP in orange, prenyl chain in yellow, and the two fluorine atoms in white. The side chain of N219 has been included to aide with orientation.

**Fig. 4.**
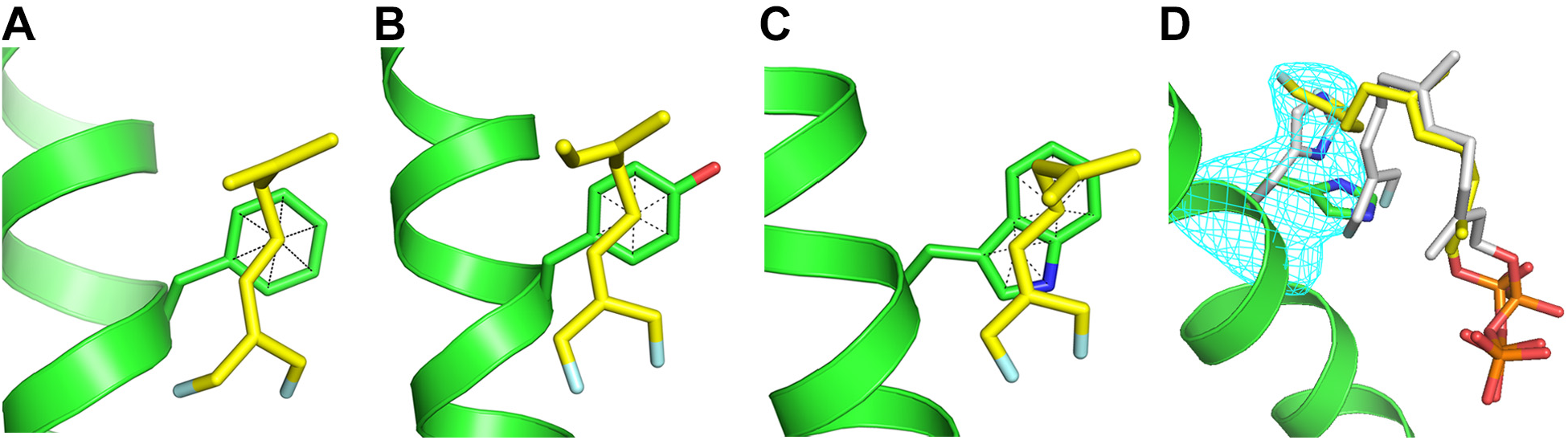
Structure of the active site surrounding residue 76 in WT and three mutants of PS. *A*, WT; *B*, F76Y; *C*, F76W; and D, F76H. The electron density (blue mesh) for F76H shown in panel D is a Polder map contoured at 3 σ and indicates the presence of at least two rotamer conformations for the 76 side chain, with one (30%) corresponding to the orientation of Phe in the WT protein (grey). Electron density was observed for DFFPP only in an extended conformation (yellow) corresponding to the new rotamer conformation of the H76 side chain (green). The grey conformation, corresponding to the ligand in WT, was modelled in to avoid steric conflict with the grey His76 rotamer.

**Table 1.**
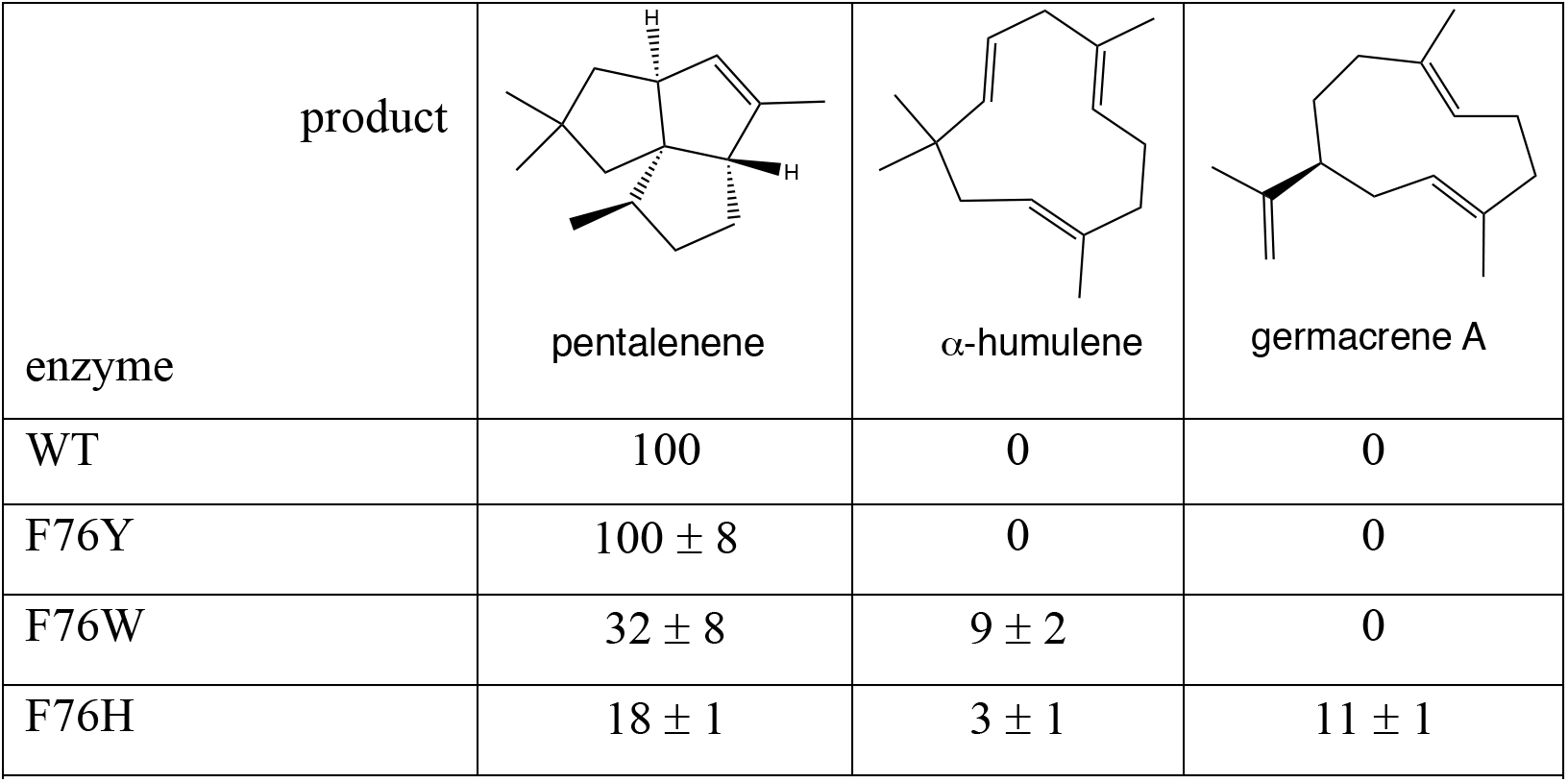
Activity and product distribution of WT and mutant PS. All values were determined by GC/MS, expressed relative to the amount of pentalenene produced by the WT enzyme (i.e., 100). Assays (overnight) were performed as described in Methods. Values are averages of two independent experiments with indicated range.

| product |  |  |  |
| --- | --- | --- | --- |
| enzyme | pentalenene | α-humulene | germacrene A |
| WT | 100 | 0 | 0 |
| F76Y | 100 ± 8 | 0 | 0 |
| F76W | 32 ± 8 | 9 ± 2 | 0 |
| F76H | 18 ± 1 | 3 ± 1 | 11 ± 1 |

To further probe the role of F76 in guiding reactivity of the C10,C11-π electrons to favor an anti-Markovnikov product, we constructed several PS mutants with amino acid substitutions at position 76. The activities and product distributions for mutants in which various aromatic amino acids replace F76 are shown in Table 1 (overnight incubations performed to examine product distributions), Figs. S3-S5 (GC-MS identification of the products), and Fig. S6 (activities under initial rate conditions). The F76A mutant, as reported by others, expresses well but is poorly soluble and not active.^*12*^ For these reasons, the Ala mutant was not pursued here. In contrast to F76A, all mutants with aromatic amino acids at position 76 display significant activity. F76Y displays near WT activity, as is expected from the calculated electrostatic potentials and gas phase Na^+^-ion binding energies for phenol versus benzene.^*26, 27*^ In addition, the Y76 side chain is found in an almost identical position in the structure of the DFFPP-bound mutant as it is in the WT protein (Fig. 4B).

The F76W mutant intriguingly generates a significant amount of a-humulene in addition to pentalenene (Table 1). a-Humulene can be generated from humulyl cation A by abstraction of a proton from C9 (see Fig. 1 and Table 1), suggesting that F76W may stabilize a carbenium ion at C10 as well as at C9. Consistent with this suggestion, the structure of the DFFPP-bound F76W mutant clearly shows carbons C9 and C10 centered above the fused benzene and pyrrole rings of the indole side chain, respectively, such that carbocations at both positions should be stabilized in this complex, leading to the two anti-Markovnikov products pentalenene and α-humulene. In addition, the indole side chain of the tryptophan may serve as a general base to facilitate deprotonation of humulyl cation A or B to generate α-humulene.

The F76H mutant is interesting. As can be seen in Table 1, F76H produces not only the two anti-Markovnikov products, pentalenene and a-humulene, but also a significant amount of germacrene A, a product resulting from Markovnikov addition of C10 to the farnesyl cation at position C1 (Fig. 1 and Table 1). Significantly, the crystal structure of the F76H/DFFPP complex displays at least two conformations of the imidazole side chain of H76 (Fig. 4D), one of which is in a similar location to that of the benzene ring of F76 in the WT protein. While density for the prenyl chain is not well defined, the two conformations of the imidazole side chain of H76 (30% and 70% by occupancy) must be associated with two dramatically different conformations of the bound DFFPP ligand because the new conformation (70%) is sterically incompatible with the ligand conformation observed in the WT protein. Thus, the mutation causes re-sculpting of the active site such that a different conformation of the ligand is stabilized in the binding pocket. These data indicate that Phe76 plays a structural role in the ligand binding pocket in addition to its role in stabilizing development of positive charge on C9, and the multiple conformations of the H76 side chain suggests a rationale, at least in general terms, for the generation of multiple products differing in regioselectivity of the initial cyclization reaction.

## Discussion

The active sites of terpene synthases are often lined with aromatic amino acid side chains, and it has long been suspected that carbocation intermediates along the reaction coordinate are stabilized through cation-π interactions involving these amino acid residues.^*5*^ Polar residues are not well-suited for this purpose as they would be at risk of covalent modification with the highly reactive carbenium ion intermediates leading to suicide inactivation of the enzymes, as has been observed for a number of different terpene synthases by Noel and coworkers.^*32*^ The presumed involvement of cation-π interactions in terpene synthase reactions has been supported by numerous studies showing that mutation of aromatic amino acid residues in terpene synthase active sites severely disrupts function of the enzymes, but the enzyme active site contour is often defined in large part by the steric bulk of aromatic amino acids, and it can be difficult to sort the effects of mutation on electrostatics from the role played by side chains in stabilizing the reactive conformation of the flexible isoprenoid substrate. In addition, with notable exceptions,^*33, 34*^ the mutagenesis data are not always supported by structural studies linking a particular aromatic residue to a carbocation intermediate in the reaction coordinate, which is crucial to understanding the mechanism by which terpene synthases guide regioselectivity in the generation of specific products from substrates which have the potential to produce a large number of different products.

An important example demonstrating the feasibility of cation-π interactions in terpene synthases was provided in a study by Christianson and coworkers in which a crystal structure of the enzyme epi-isozizaene synthase in complex with 3 Mg^2+^ ions, inorganic pyrophosphate, and the benzyltriethylammonium cation showed the quaternary ammonium ion surrounded by three phenylalanine side chains in the active site about 5 Å from the center of each aromatic side chain.^*38*^ While this example provides direct demonstration of cation-π interactions in the active site of a terpene synthase, a comparable example with a substrate analog does not exist, and this is especially true for terpene synthases catalyzing reactions in which anti-Markovnikov additions are a part of the overall reaction mechanism. The results presented here provide compelling evidence for the involvement of Phe76 in controlling regioselectivity for developing carbenium ions in the PS reaction through stabilizing cation-π interactions with the benzene ring of the side chain. Following a soak of the apo-PS crystal, the DFFPP substrate analog was found in the active site with C9 positioned 3.5 Å above the center of the benzene ring of the side chain, perfectly placed for stabilization of a developing carbenium ion through cation-π interaction with the aromatic ring on one face and mainchain carbonyl dipole of I177 and neighboring intramolecular C6,C7-double bond on the opposite face. The fact that C10 is found outside of the perimeter of the benzene ring suggests that positive charge may not significantly develop on C10 in the reaction coordinate, and the 1,2-hydride shift transferring charge to C9 may coincide with initial attack of C11 on C1 in a concerted reaction mechanism. This proposal is supported by the crystal structure and product profile of the F76W mutant in which the indole ring of the side chain is in position to support development of positive charge on C10, and the product profile includes a-humulene, which likely results from deprotonation of a humulyl cation A intermediate (Fig. 1) in the mutant that may not exist in the WT protein.

Finally, we address how conserved the F76 residue is in terpene synthases. The search is restricted to *Streptomyces*, the largest group of bacterial sesquiterpene synthases. There is a rough correlation with enzymes catalyzing reactions that begin with anti-Markovnikov attack and development of positive charge on C9 (Figs. 5 & 6). Pentalenene synthase, cucumene synthase,^*36*^ isohirsut-4-ene synthase,^*37*^ and (+)-isoafricanol synthase^*35*^ all begin with attack of C11 on C1, followed by a 1,2 hydride shift resulting in a carbenium ion intermediate with positive charge on C9, and all have either a Phe or Tyr residue at position 76 (pentalenene synthase numbering). In contrast, a-eudesmol synthase,^*39*^ a-amorphene synthase,^*39*^ selina-4(15),7(11)-diene synthase,^*31*^ daucadiene synthase,^*40*^ (+)-T-muurolol synthase,^*41*^ germacradiene-11-ol synthase,^*39*^ (-)-δ-cadinene synthase,^*41*^ and (+)-epicubenol synthase^*42*^ all have a nonaromatic amino acid residue at position 76 and none catalyze reactions that are initiated with anti-Markovnikov addition reactions. However, the correlation is not absolute. Epi-isozizaene synthase,^*43*^ (-)-epi-a-bisabolol synthase,^*44*^ avermitilol synthase,^*45*^ and (-)-germacradien-4-ol synthase^*44*^ all have an aromatic amino acid side chain at position 76, but none catalyze a reaction beginning with an anti-Markovnikov attack, and africanene synthase^*40*^ does not have an aromatic residue at position 76, but does catalyze a reaction beginning with an anti-Markovnikov attack and development of positive charge on C9.

**Figure 5.**
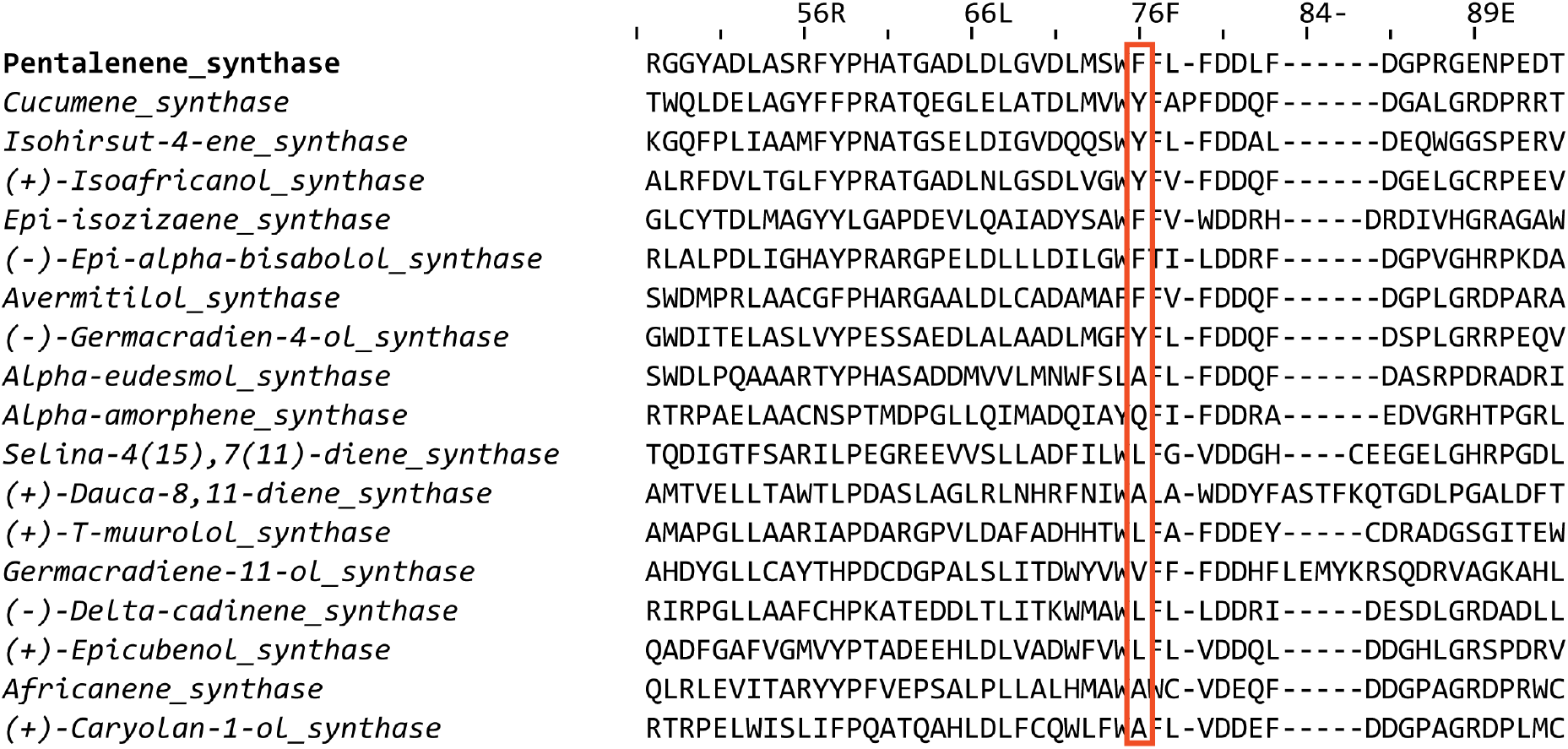
Multiple sequence alignment of sesquiterpene synthases from *Streptomyces sp*. Full length sequences for pentalenene synthase (AAA19131.1), cucumene synthase (B5GLM7), isohirsut-4-ene synthase (slt18_1880), (+)-isoafricanol synthase (from *S. malaysiensis*), epi-isozizaene synthase (Q9K499), epi-a-bisabolol synthase (AB621339), avermitilol synthase (Q82RR7), (-)-germacradien-4-ol synthase (AB621338), α-eudesmol synthase (SCNRRL3882_07544), a-amorphene (SCNRRL3882_07041), selina-4(15),7(11)-diene synthase (B5HDJ6), dauca-8,11-diene synthase (SVEN_0552), (+)-T-muurolol synthase (B5GW45), germacrenediene-11-ol-synthase (SCNRRL3882_01776), (-)-δ-cadinene synthase (B5GS26), (+)-epicubenol synthase (B1W477), africanene synthase (SCLAV_p0985), and (+)-caryolan-1-ol synthase (B1W019.1) were aligned. The sequence for pentalenene synthase was used as reference for preparing the figure, with the column containing F76 in pentalenene synthase enclosed in a red box. Sequences were aligned using MUSCLE^*1, 2*^ and visualized with Jalview 2.^*3*^

**Figure 6.**
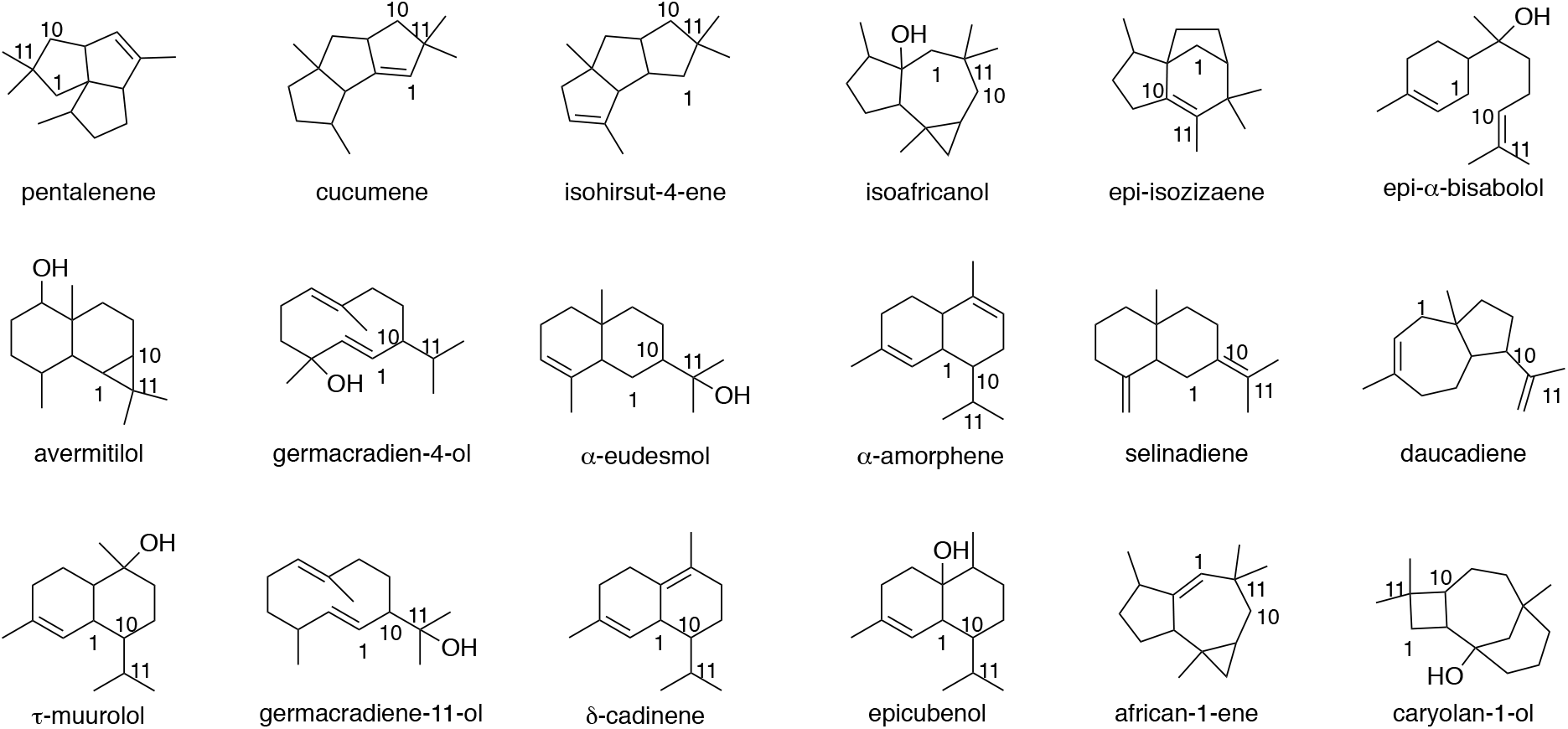
Chemical structures of compounds produced by the enzymes in the multiple sequence alignment.

(+)-Caryolan-1-ol synthase (CS) is an interesting case (Fig. 7). CS catalyzes formation of the tertiary alcohol caryolanol from FPP. As with PS, the reaction scheme begins with anti-Markovnikov attack of C11 on C1 of the farnesyl cation, but in this case the secondary carbocation remains localized to C10 such that subsequent attack of C2 onto C10 forms a fused four-membered cyclobutane ring instead of the five-membered ring of the PS reaction.

**Fig. 7.**
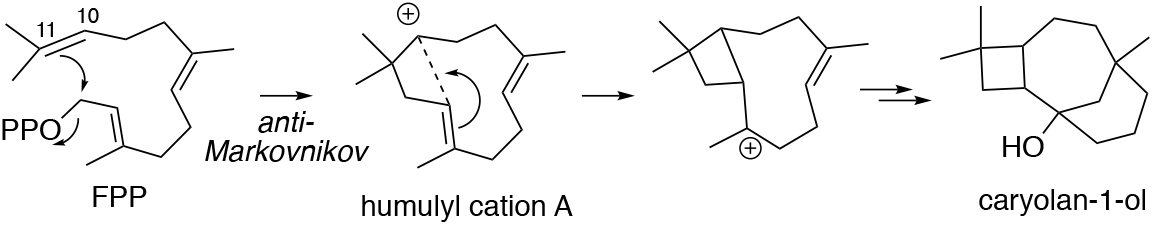
Reaction scheme for (+)-caryolan-1-ol synthase.

Our interest in this reaction stems from its initial similarity to the PS reaction coupled with subsequent differences to generate a distinct product. In addition, the enzyme from *Streptomyces griseus*, which shares 29% amino acid identity with PS, notably, has an Ala in the same position as F76 in PS (Fig. 5). Given that the two proteins share only 29% sequence identity, it seems unlikely that substitution of Ala with Phe at this position would create a pentalenene synthase (and we have confirmed that suspicion), but we also guess that exploration of the CS reaction will deepen significantly our understanding of how terpene synthases control regioselectivity in the cyclization reactions.

## Methods

### Synthesis of DMAPP, IPP, and *E,E*-FPP

The synthesis and purification of DMAPP, IPP, and *E,E*-FPP from the corresponding allylic alcohols (Sigma-Aldrich, MO) was performed as previously described for substrates and substrate analogs of the enzyme (+)-limonene synthase.^*21, 46, 47*^ The purity of the diphosphate products was assessed by proton, carbon, and phosphorus NMR. NMR spectra were recorded on a Varian 400-MR spectrometer (9.4 T, 400 MHz) in D_2_O adjusted to pD ~8.4 with ND_4_OD. ^1^H and ^13^C chemical shifts are reported in parts per million (ppm) downfield from TMSP (trimethylsilyl propionic acid). ^31^P chemical shifts are reported in ppm relative to 85% *o*-phosphoric acid. J coupling constants are reported in units of frequency (hertz) with multiplicities listed as s (singlet), d (doublet), dd (doublet of doublets), t (triplet), q (quartet), m (multiplet), br (broad), and app (apparent).

Dimethylallyl diphosphate (DMAPP): ^1^H NMR (400 MHz, D_2_O/ND_4_OD) δ_H_ 1.73 (3 H, s, CH3), 1.77 (3 H, s, CH3), 4.46 (2 H, app t, J = 6.8 Hz at C1), 5.46 (1 H, t, J = 7.4 Hz at C2); ^13^C{^1^H} NMR (100 MHz, D_2_O/ND_4_OD) δ_C_ 20.13, 27.79, 65.37 (d, J_CP_ = 5.3 Hz), 122.72 (d, J_CP_ = 8.5 Hz), 142.71; ^31^P{^1^H} NMR (162 MHz, D_2_O/ND_4_OD) δ_P_ −6.76 (d, J_PP_ = 22.1 Hz, P1), – 10.46 (dt, J_PP_ = 22.1 Hz, J_PH_ = 6.8 Hz P2).

Isopentenyl diphosphate (IPP): ^1^H NMR (400 MHz, D_2_O/ND_4_OD) δ_H_ 1.78 (3 H, s, CH3 at C4), 2.40 (2 H, t, J = 6.4 Hz at C2), 4.07 (2 H, app q, J = 6.7 Hz at C1), 4.85 (1H, s, H at C5), 4.87 (1H, s, H at C5); ^13^C{^1^H} NMR (100 MHz, D_2_O/ND_4_OD) δ_C_ 24.58, 40.77 (d, J_CP_ = 7.6 Hz), 66.90 (d, J_CP_ = 5.4 Hz), 114.32, 146.85; ^31^P{^1^H} NMR (162 MHz, D_2_O/ND_4_OD) δ_P_ −6.60 (d, J_PP_ = 22.0 Hz, P1), −10.60 (d, J_PP_ = 22.0 Hz, J_PH_ = 6.6 Hz, P2).

*E,E*-Farnesyl Diphosphate (FPP): ^1^H NMR (400 MHz, D_2_O/ND_4_OD) δ_H_ 1.63 (6H, two s, CH3), 1.70 (3 H, s, CH3), 1.73 (3 H, s, CH3), 2.00-2.23 (8 H, m, H at C4, C5, C8, C9), 4.48 (2 H, app t, J = 6.5 Hz, H at C1), 5.16-5.26 (2 H, m, H at C6, and C10), 5.48 (1 H, t, J = 7.2 Hz at C2); ^13^C{^1^H} NMR (100 MHz, D_2_O/ND_4_OD) δ_C_ 18.11, 18.51, 19.83, 27.71, 28.51, 28.63, 41.65, 41.71, 65.33 (d, J_CP_ = 5.2 Hz), 122.82 (d, J_CP_ = 8.5 Hz), 127.17, 127.33, 136.40, 139.57, 145.61; ^31^P{^1^H} NMR (162 MHz, D_2_O/ND_4_OD) δ_P_ −5.76 (d, J_PP_ = 22.2 Hz, P1), −9.59 (d, J_PP_ = 22.2 Hz, J_PH_ = 6.5 Hz, P2).

### Preparation of FPPS and PS

Genes for farnesyl pyrophosphate synthase (*fpps*) from *T. cruzi* (genbank AF312690) and pentalenene synthase (*ps*) from *Streptomyces sp. UC5319* (GenBank UO5213) were codon optimized for expression in *E. coli* and purchased from Synbio Technologies (New Jersey). The genes were in a pET-28a (+) vector as a NcoI (5’-end) and EcoRI (3’-end) cartridge with a glycine codon immediately after the start methionine and followed by a His6 tag and tobacco etch virus (TEV) protease cut site.

The plasmids were used to transform BL21 (DE3) competent cells, and colonies selected for growth on LB/agar plates containing 50 μg/mL kanamycin. A single colony was used to inoculate 10-15 mL of LB containing 50 μg/mL kanamycin, and then incubated overnight at 37 °C with shaking at 220 rpm. The overnight culture was used to inoculate 1 L of LB containing 50 μg/mL kanamycin. General procedures for purification of the proteins were performed as previously described for (+)-limonene synthase.^*47*^ Procedures specific for each protein were essentially as previously described.^*9, 48*^

FPPS. Inoculated cultures were grown with shaking at 37 °C to OD_600_ ≈ 0.6 and induced with 1 mM IPTG, as described previously.^*48*^ Harvested cells were re-suspended in 20 mM Tris buffer, pH 7.9, containing 500 mM NaCl and 5 mM DTT (Buffer A), sonicated, and clarified by centrifugation. The resulting lysate was loaded at a rate of 2 mL/min onto a prepacked 5 mL HiTrapFF Ni-Sepharose column (GE Healthcare Life Sciences, MA) that had been equilibrated with Buffer A. The column was washed with 30 mL of 20 mM imidazole in Buffer A and then eluted with a linear gradient of 20-250 mM imidazole in Buffer A. Fractions containing FPPS were pooled, the imidazole removed by repeated concentration using Amicon Ultra – 15 filtration units with a 10 kDa cutoff followed by dilution with Buffer A, and the sample aliquoted (100 μM), then flash frozen with liquid nitrogen, and stored at −80°C until further use.

PS.^*9*^ 1 L cultures were grown at 37 °C with shaking at 220 rpm to an OD_600_ ~0.4-0.6, then moved to a pre-cooled shaking incubation at 16 °C, and induced with 0.5 mM IPTG after 1 hr, and incubated overnight. Cells were harvested by centrifugation and resuspended in 50 mM bistris propane buffer, pH 7.5, containing 150 mM KCl, and 10 mM MgCl2 (Buffer B), sonicated, and the supernatant fraction clarified by centrifugation. The resulting lysate was loaded at a rate of 2 mL/min onto a prepacked 5 mL HiTrapFF Ni-Sepharose column (GE Healthcare Life Sciences, MA) that had been equilibrated in Buffer B. The column was then washed with ~30 mL of Buffer B containing 20 mM imidazole, and the protein eluted with a linear gradient of 0.02-1.0 M imidazole in Buffer B. For PS mutants, which tended to be less stable in the presence of high imidazole concentrations, fractions were collected in 50-mL Falcon tubes containing 10 mL of Buffer B to dilute the imidazole. Eluted protein was buffer exchanged into low salt buffer (50 mM Tris base, 5 mM DTT, 10% glycerol at pH 8.2) and loaded onto a 5 mL prepacked Q-sepharose ion exchange column (GE Healthcare Life Sciences, MA) at 1 mL/min. Protein was eluted with a linear NaCl gradient ranging from 0 to 1M. Fractions containing PS were pooled, the imidazole removed by repeated concentration using Amicon Ultra – 15 filtration units with a 10 kDa cutoff followed by dilution with Buffer B, and the sample aliquoted (100 μM), then flash frozen with liquid nitrogen, and stored at −80 °C until further use.

### Preparation and Characterization of 12,13-Difluorofarnesyl Diphosphate (DFFPP)

DFFPP was prepared enzymatically from DFGPP (Fig. 2). 10 mM 8,9-DFGPP,^*21, 22*^ 10 mM IPP, and 10 mM MgCl2 were incubated with 5 μM FPPS in 450 μL of 20 mM Tris-buffer, pH 7.9, containing 500 mM NaCl overnight at RT, and the crude reaction mixture then stored at −20 °C. The thawed sample was filtered by centrifugation through an Amicon Ultra filter with 10 kDa molecular mass cutoff, and the flow-through fraction collected and lyophilized to dryness. After re-suspension of the lyophilized powder in 500 μL of D_2_O, and recording of a ^1^H NMR spectrum, the presence of 3 vinyl protons matched a previously reported spectrum^*22*^ of 12,13-DFFPP and indicated that the enzymatic elongation of 8,9-DFGPP to 12,13-DFFPP had been successful. The sample was then lyophilized, re-suspended in 50 μL H2O, and stored at −20 °C until use. To determine if PS was unreactive with DFFPP, 1 μM FPPS, 1 μM PS, 200 μM DFGPP, and 200 μM IPP was prepared in a final volume of 1 mL of buffer B and overlaid with 1 mL of hexanes and incubated overnight at RT (Fig. S8d). A control reaction containing 200 μM DMAPP in lieu of 8,9 DFGPP and 400 μM IPP was prepared in the same fashion (Fig. S7a). To ensure FPPS converts 8,9 DFGPP and IPP into DFFPP, 1 μM FPPS was mixed with 200 μM 8,9 DFGPP, 200 μM IPP in a total volume of 1 mL of buffer B and incubated overnight at RT. 100 μL of 1N HCl was then added to the reaction and incubated at RT for 2 hr to convert the diphosphates into the corresponding alcohols. The reaction mixtures were then neutralized with 100 μL of 1 N NaOH before analyzing the sample by GC-MS (Fig. S7e & f). A control reaction containing 1 μM FPPS, 200 μM DMAPP, and 400 μM IPP was also prepared and treated with acid, extracted with 1 mL hexanes, and the isoprenoid alcohols analyzed by GC-MS (Fig. S7b & c).

### Enzyme Activity

Enzyme assays were performed using the discontinuous single-vial method described by O’Maille et al.,^*49*^ with the exception that hexanes were used for the organic layer rather than ethyl acetate. Hexane extractable terpene products were identified and quantified using GC-MS (Agilent Technologies 7890A GC System coupled with a 5975C VL MSD with a triple-axis detector) as previously described by us and others.^*47, 50*^ Pulsed-splitless injection was used to inject 5 μL samples onto an HP-5ms (5%-phenyl)-methylpolysiloxane capillary GC column (Agilent Technologies, CA; 30 m × 250 μm × 0.25 μm) at an inlet temperature of 220 °C, transfer temperature of 240 °C, and run at constant pressure using helium as the carrier gas. Samples were initially held at an oven temperature of 50 °C for 1 min, followed by a linear temperature gradient of 10 °C/min to 220 °C, which was then held for 10 min.

The assays were run under two different conditions: 1. Overnight reactions to determine product distribution; and 2. Initial-rate reactions to determine kcat for the major product. The concentrations of the different products were estimated on the basis of a standard curve for α-humulene.

Overnight reactions. 1 μM of PS (either WT, F76Y, F76W, F76H, or F76S) was mixed with 200 μM of *E,E*-FPP in a total reaction volume of 1 mL buffer B. The solution was overlaid with 1 mL hexanes (Sigma-Aldrich, MO) and incubated overnight at RT. The solutions were vortexed to stop the reaction and extract products into the organic layer which was analyzed by GC-MS. Purchased standards of α-elemene (Cayman Chemical, MI) and a-humulene (Sigma-Aldrich, MO) were used to determine identity of major, non-pentalenene peaks from reactions of the PS mutants, where product formation was followed in scan mode.

Initial rates. Initial rates were determined under Vmax conditions. Enzyme (10 nM for PS WT and F76Y; 100 nM for F76W, and F76H) was mixed with 50 μM *E,E*-FPP (75 μM for F76W) in 1 mL buffer B, overlaid with 1 mL hexanes, and incubated at RT for various times before vortexing to stop the reaction and extract products into the organic layer for analysis by GC-MS as described above, where product formation was followed in select ion monitoring (SIM) mode, specifically set to monitor a mass at 204.1.

### Crystallization

Crystallization trials were performed by sitting drop vapor diffusion at room temperature using Hampton (Hampton Research, CA) and Jena Bioscience (Jena Bioscience, Jena, Germany) sparse matrix crystallization screens. Drops were set with a Phoenix robot (Art Robbins Instruments, CA) by mixing protein (7.5 mg/mL) in 50 mM Bis-Tris propane, pH 7.5, 150 mM KCl and 10 mM MgCl2 with crystallization mother liquor in a 1:1 ratio. Crystals of WT-Mg appeared after about 3 weeks in 8-14% w/v PEG4000, 100 mM HEPES, pH 6.5-7.5, 200 mM MgCl2, 15% ethylene glycol and 10% 2-propanol at 20 °C while WT and mutant crystals formed in 0.8-1.2 M sodium tartrate, 100 mM Tris, pH 7.5-9, and 5 mM DTT.

### Data Collection, Processing, and Refinement

WT-Mg crystals were soaked in reservoir solution containing 15% glycerol as cryoprotectant before flash freezing in liquid nitrogen. To generate the DFFPP complexes, WT and mutant crystals were soaked with reservoir solution containing 1 mM DFFPP, 10 mM MgCl_2_, and 15% glycerol for 1 hr at room temperature before flash-freezing in liquid nitrogen. Diffraction data were collected at 100 K with beam line 5.0.2 at the Advanced Light Source (Lawrence Berkeley National Laboratory, Berkeley, CA) using a Pilatus3 X 6M detector (DECTRIS Ltd., Switzerland). Data sets were integrated using XDS^*51*^ and scaled using SCALA version 3.3^*52*^ from the CCP4 software suite version 7.0.^*53, 54*^ All diffraction data sets, except WT-Mg, were processed in space group *P*6_3_. WT-Mg data were processed in *P*2_1_. Complete data collection statistics are listed in Table 1.

The WT structure was solved by molecular replacement using PHASER version 2.8^*55*^ with molecule A of the published structure for PS (PDB entry 1PS1) as a search model. All other structures were solved using the final refined model of WT as a search model. The coordinates of WT have not been deposited, as the structure is essentially identical to that of PDB entry 1PS1. The molecular replacement solutions contain two molecules in the asymmetric unit. Rigid body refinement followed by positional and B-factor refinement was carried out using phenix.refine^*56*^ from the PHENIX software suite version 1.16.^*57*^ Simulated annealing was included in earlier refinements to minimize the initial model bias. Manual model building was done using COOT version 0.8.^*58*^ After initial refinement, difference Fourier electron density was observed for the analog in the active site of the complex protein structures. The ligand was modeled using Jligand version 1.0 from CCP4 software suite version 7.0 and the generated coordinates and restraints were used for further refinements. Water molecules were included in the final refinement after satisfying the criteria of 3 σ *F_o_ – F_c_* and 1 σ *2F_o_ – F_c_*. Several iterative cycles of refinement were carried out before final submission of data. Data collection and final refinement statistics are given in Table 2. Data sets for the WT-Mg (PDB entry 6WKC), WT-DFFPP (PDB entry 6WKD), F76Y (PDB entry 6WKE), F76Y-DFFPP (PDB entry 6WKF), F76W (PDB entry 6WKG) and F76W-DFFPP (PDB entry 6WKH), F76H (PDB entry 6WKI), and F76H-DFFPP (PDB entry 6WKJ) have been submitted to the Protein Data Bank. All crystal structure figures in this paper were prepared using PyMol version 2.3 (Schrödinger LLC, Portland, OR).

**Table 2.** Crystallographic Data Collection and Refinement Statistics.

|  | WT-MGapo | WT-DFFPP | F76Y-apo | F76Y-DFFPP | F76W-apo | F76W-DFFPP | F76H-apo | F67H-DFFPP |
| --- | --- | --- | --- | --- | --- | --- | --- | --- |
| *PDB ID* | 6WKC | 6WKD | 6WKE | 6WKF | 6WKG | 6WKH | 6WKI | 6WKJ |
| *Data collection statistics* |  |  |  |  |  |  |  |  |
| Space group | $P2_1$ | $P6_3$ | $P6_3$ | $P6_3$ | $P6_3$ | $P6_3$ | $P6_3$ | $P6_3$ |
| Resolution<br>range (Å) | 20 - 1.65 | 20 - 2.20 | 20 - 2.40 | 20 - 2.50 | 20.0 - 2.30 | 20.0 - 2.55 | 20 - 2.35 | 20.0 - 2.30 |
| Highest<br>resolution shell<br>(Å) | 1.74 - 1.65 | 2.32 - 2.20 | 2.53 - 2.40 | 2.64 - 2.50 | 2.42 - 2.30 | 2.69 - 2.55 | 2.48 - 2.35 | 2.42 - 2.30 |
| Unit cell<br>parameters (Å) | a = 60.7,<br>b = 73.8,<br>c = 81.2;<br>$\alpha = \gamma = 90$,<br>$\beta = 100.2$ | a = 182.5,<br>b = 182.5,<br>c = 56.2;<br>$\alpha = \beta = 90$,<br>$\gamma = 120$ | a = 181.8,<br>b = 181.8,<br>c = 56.4;<br>$\alpha = \beta = 90$,<br>$\gamma = 120$ | a = 183.3,<br>b = 183.3,<br>c = 56.7;<br>$\alpha = \beta = 90$,<br>$\gamma = 120$ | a = 183.2,<br>b = 183.2,<br>c = 56.5;<br>$\alpha = \beta = 90$,<br>$\gamma = 120$ | a = 185.0,<br>b = 185.0,<br>c = 56.9;<br>$\alpha = \beta = 90$,<br>$\gamma = 120$ | a = 181.5,<br>b = 181.5,<br>c = 56.5;<br>$\alpha = \beta = 90$,<br>$\gamma = 120$ | a = 182.0,<br>b = 182.0,<br>c = 56.4;<br>$\alpha = \beta = 90$,<br>$\gamma = 120$ |
| Total<br>reflections | 566358 | 1114810 | 853949 | 755975 | 951335 | 726238 | 911638 | 952163 |
| Unique<br>reflections | 83734 | 54634 | 41985 | 38010 | 48511 | 36655 | 44617 | 47794 |
| Completeness<br>%<sup>a</sup> | 98.9 (98.0) | 99.9 (100) | 99.8 (100) | 99.8 (100) | 99.8 (100) | 99.8 (100) | 99.8 (100) | 99.8 (100) |
| R<sub>merge</sub> %<sup>a</sup> | 9.4 (91) | 11.4 (172) | 12.8 (152) | 16.5 (112) | 13.6 (146) | 13.7 (167) | 11.1 (179) | 10.9 (150) |
| I/σ (I)<sup>a</sup> | 14.9 (2.0) | 22.2 (2.2) | 21.6 (2.4) | 9.4 (2.3) | 16.9 (2.5) | 15.9 (2.1) | 23.5 (2.0) | 20.0 (2.3) |
| Redundancy<sup>a</sup> | 6.8 (6.7) | 20.4 (21.0) | 20.3 (21.1) | 19.9 (20.5) | 19.6 (20.2) | 19.8 (20.2) | 20.4 (20.5) | 19.9 (20.3) |
| *Refinement statistics* |  |  |  |  |  |  |  |  |
| Resolution<br>range (Å) | 20 - 1.65 | 20 - 2.20 | 20 - 2.4 | 20.0 - 2.5 | 20 - 2.30 | 20 - 2.55 | 20 - 2.35 | 20 - 2.3 |
| No. of<br>reflections<br>used | 83593 | 54616 | 41971 | 37825 | 48499 | 36638 | 44601 | 47769 |
| R<sub>cryst</sub> % | 16.7 | 20.1 | 19.8 | 20.8 | 21.6 | 22.5 | 22.0 | 21.1 |
| R<sub>free</sub> % | 19.3 | 21.9 | 23.2 | 23.5 | 23.7 | 25.6 | 24.1 | 24.6 |
| Protein atoms | 4842 | 4611 | 4494 | 4430 | 4455 | 4377 | 4493 | 4355 |
| Ligand atoms | 0 | 26 | 0 | 26 | 0 | 26 | 0 | 26 |
| Metal atoms | 4 | 2 | 0 | 0 | 0 | 0 | 0 | 0 |
| Water molecules | 1171 | 363 | 234 | 97 | 304 | 101 | 172 | 212 |
| r.m.s.d. in bond lengths (Å) | 0.005 | 0.003 | 0.003 | 0.003 | 0.002 | 0.002 | 0.002 | 0.004 |
| r.m.s.d. in bond angles (°) | 0.8 | 0.5 | 0.5 | 0.5 | 0.4 | 0.4 | 0.4 | 0.6 |

<sup>a</sup> Highest resolution shell values are given in parentheses.

## Acknowledgements

We are grateful to the staff at the Advanced Light Source-Berkeley Center for Structural Biology for their assistance in X-ray data collection. The Advanced Light Source is funded by the Director, Office of Science, Office of Basic Energy Sciences, of the United States Department of Energy under contract DE-AC02-05CH11231. The Berkeley Center for Structural Biology is supported in part by grants from the NIGMS, National Institutes of Health. We would like to thank Prof. Barry Snider for helpful discussions.

## Author contributions

J.O.M., R.P.K., I.J.K., and D.D.O. were involved with conceptualization and design of the overall project. A.C.M., M.P., and J.O.M were involved with most of the experiments. R.P.K. was responsible for x-ray crystallography. J.O.M., R.P.K., I.J.K., and D.D.O. wrote the manuscript.

