## Supplementary material for "Mechanism Underlying Anti-Markovnikov Addition in the Reaction of Pentalenene Synthase": Fig S1-7

### AUTHOR INFORMATION

† These authors contributed equally to this work

**Corresponding Authors**

*D.D.O., Department of Biochemistry, Brandeis University, 415 South St., Waltham, MA 02454. Telephone: 781-736-2322. Fax: 781-736-8487..

*I.J.K., Department of Chemistry, Brandeis University, 415 South St., Waltham, MA 02454. Telephone: 781-736-2574. Fax: 781-736-2516..

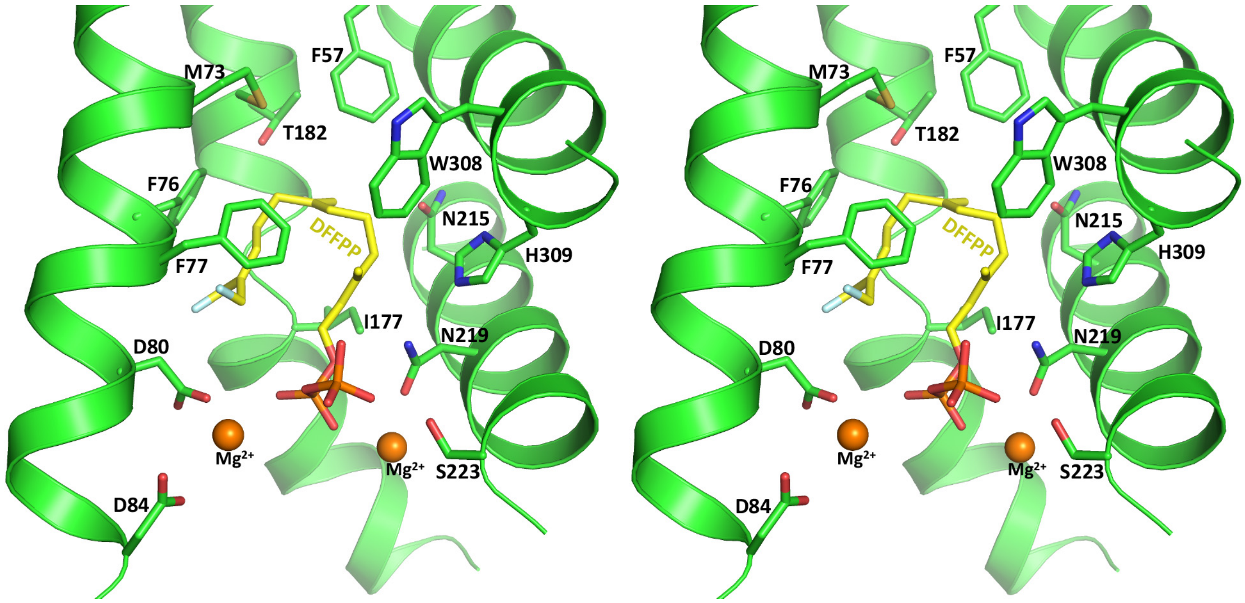


**Fig. S1. Stereo view of DFFPP ligand and metal ions in the active site of PS.**

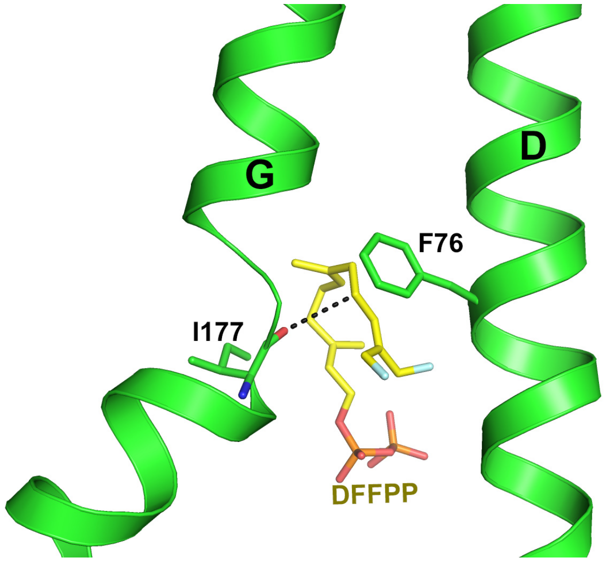


**Fig. S2. Position of the I177 mainchain carbonyl with respect to C9 of the DFFPP substrate analog.**

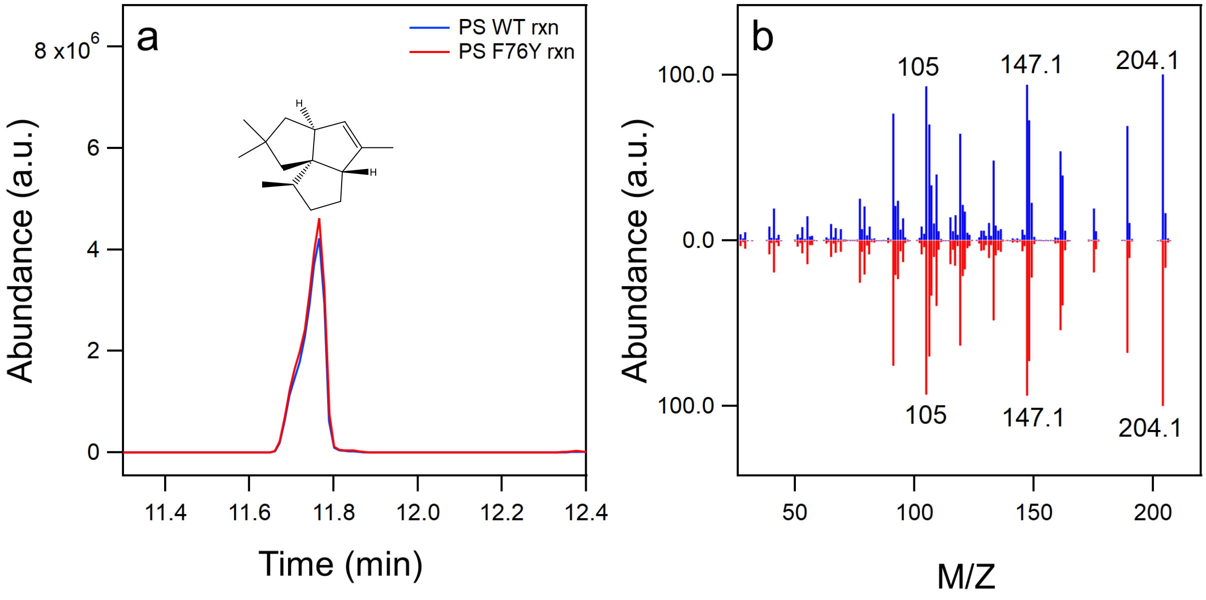


**Figure S3. Product distribution profile for PS WT and PS F76Y.** (a) Enzymatic reactions containing 1 μM enzyme and 200 μM *E,E*-FPP were incubated overnight. (b) Mass fragmentation pattern pentalenene peak at 11.75 minutes from WT (blue) and the F76Y mutant (red).

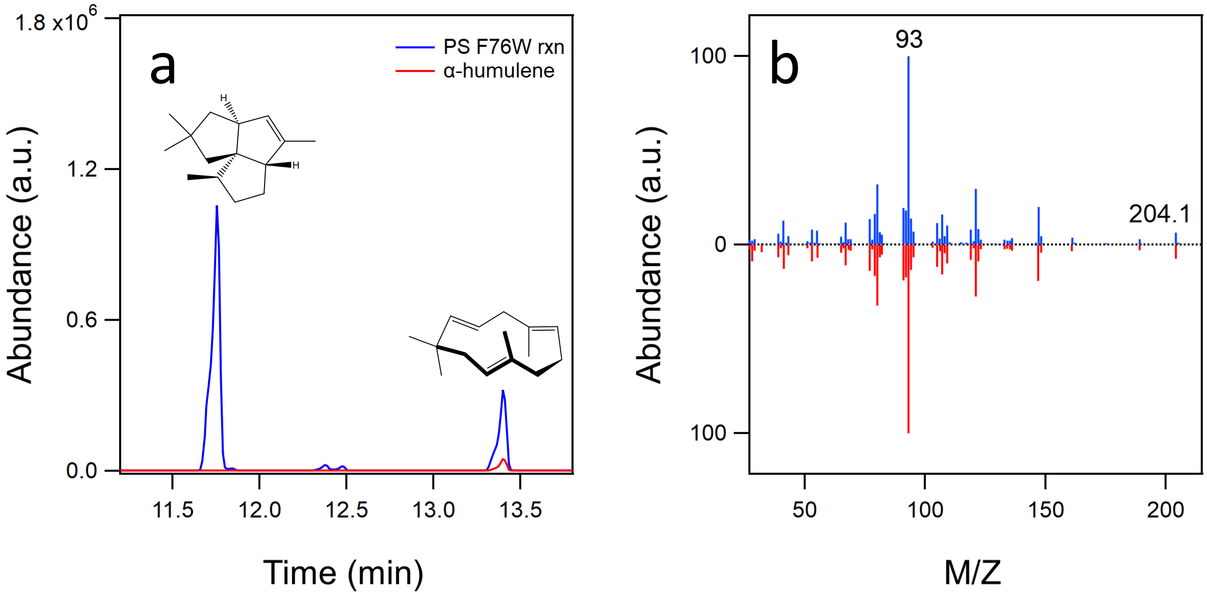


**Figure S4. Product distribution profile for PS F76W.** (a) Enzymatic reaction containing 1 μM PS F76W and 200 μM *E,E*-FPP was incubated overnight. The peak at 13.39 minutes was identified α-humulene. (b) Mass fragmentation pattern for the enzymatically-produced α-humulene (blue) and a purchased standard (red). MS data were normalized to facilitate comparison.

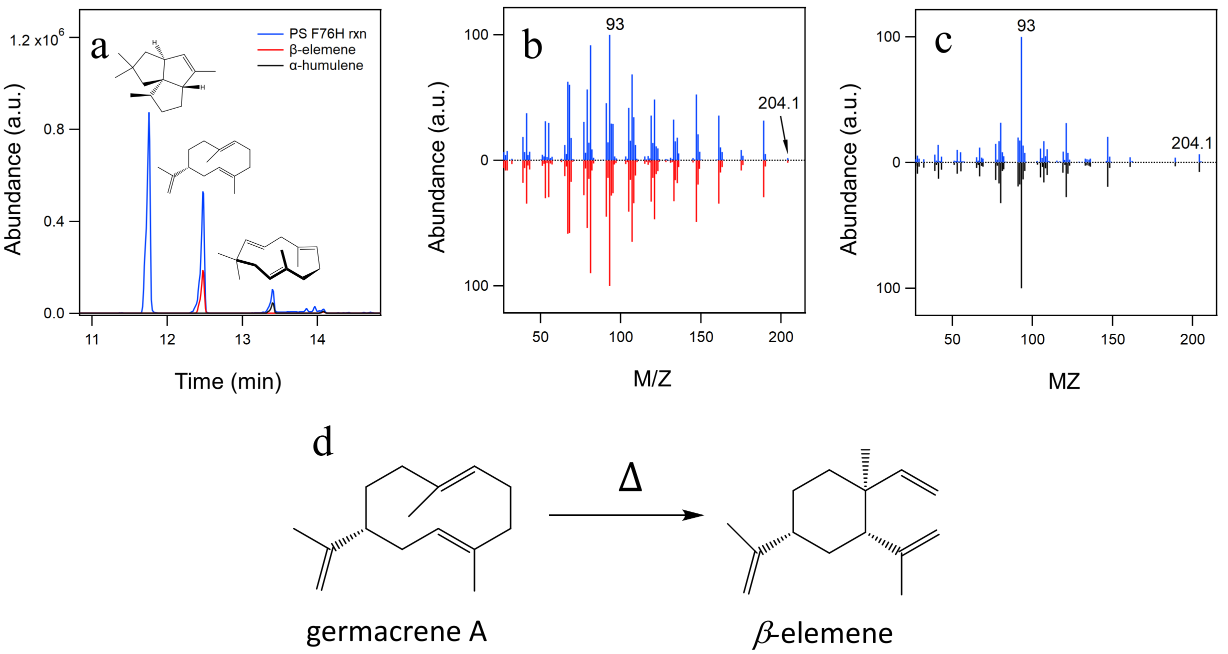


**Figure S5. Product distribution profile for PS F76H.** (a) Enzymatic reactions containing 1 µM PS F76H and 200 µM *E,E*-FPP were incubated overnight. Peaks at 12.47 and 13.39 minutes were identified as β-elemene, resulting from the cope rearrangement of germacrene A, and α- humulene, respectively. (b) Mass fragmentation pattern for enzymatically- produced germacrene A (blue) and a purchased standard of β-elemene (red). (c) Mass fragmentation pattern for enzymatically produced α-humulene (blue) and a purchased standard of α- humulene (black). (d) Scheme showing the heat driven cope rearrangement of germacrene A to β-elemene. MS data were normalized to facilitate comparison.

| | $k_{cat}$ ($s^{-1}$) | | |
|---|---|---|---|
| | Pentalenene | α-Humulene | Germacrene A |
| WT | 0.46 ± .08 | | |
| F76Y | 0.36 ± .06 | | |
| F76W | $2.1 \times 10^{-3} \pm 1 \times 10^{-4}$ | $5.9 \times 10^{-4} \pm 6 \times 10^{-5}$ | |
| F76H | $1.6 \times 10^{-3} \pm 1 \times 10^{-4}$ | $4.2 \times 10^{-3} \pm 3 \times 10^{-4}$ | $2.2 \times 10^{-2} \pm 1 \times 10^{-3}$ |

**Figure S6. $K_{cat}$ values for PS WT and F76 mutants.** All experiments were conducted under initial rate conditions as described in Methods and performed at least in triplicate.

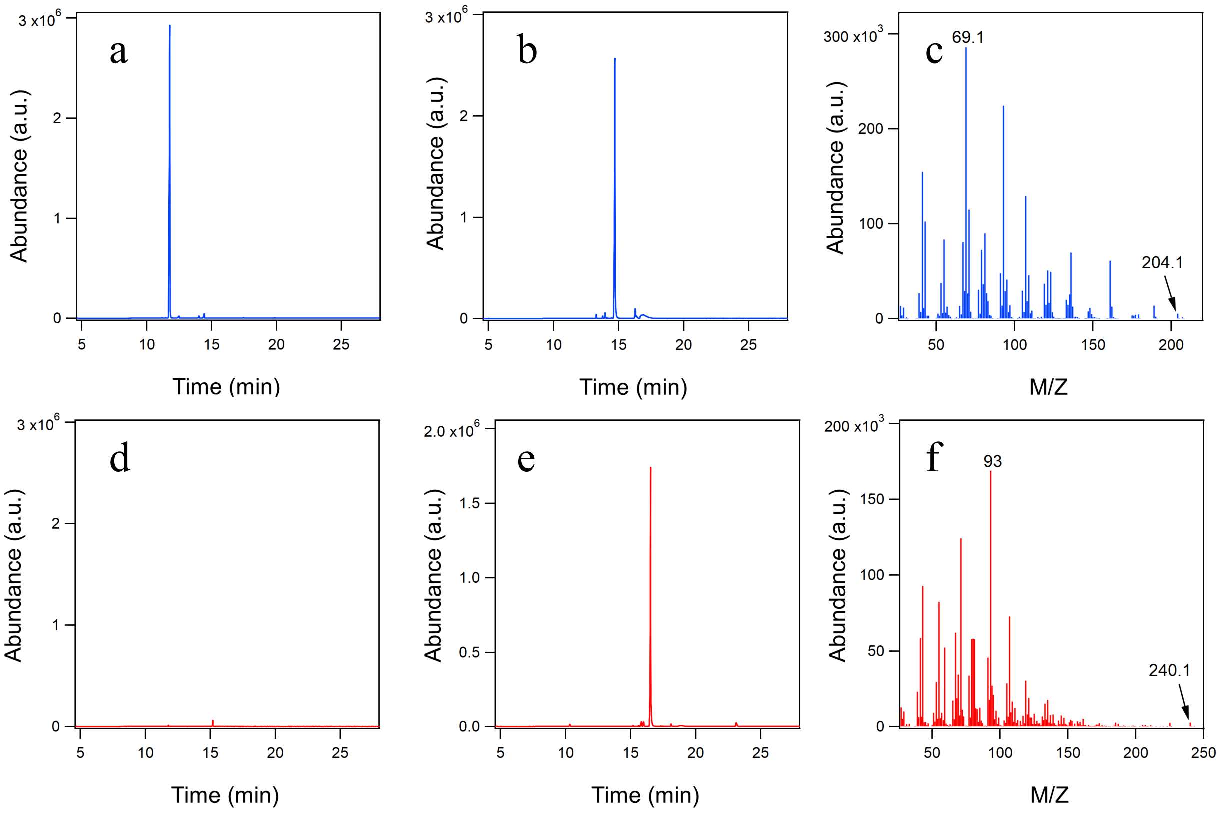


**Figure S7. Characterization of enzymatically prepared DFFPP.** (a) Product profile for reaction of 1 μM FPPS, 1 μM PS, 200 μM DMAPP, and 400 μM IPP. (b) Product profile for reaction of 1 μM FPPS, 200 μM DMAPP, and 400 μM IPP following acidification of the reaction mixture. (c) Mass fragmentation pattern for the major peak in *panel b*. (d) Product profile for reaction of 1 μM FPPS, 1 μM PS, 200 μM 8,9-DFGPP, and 200 μM IPP. (e) Product profile for reaction of 1 μM FPPS, 200 μM 8,9-DFGPP, and 200 μM IPP following acidification of the reaction mixture. (f) Mass fragmentation pattern for the major peak in *panel e*.
